# Magnetic Field Mapping and Correction for Moving OP-MEG

**DOI:** 10.1101/2021.05.25.444975

**Authors:** Stephanie Mellor, Tim M. Tierney, George C. O’Neill, Nicholas Alexander, Robert A. Seymour, Niall Holmes, José D. López, Ryan M. Hill, Elena Boto, Molly Rea, Gillian Roberts, James Leggett, Richard Bowtell, Matthew J. Brookes, Eleanor A. Maguire, Matthew C. Walker, Gareth R. Barnes

## Abstract

**Background:** Optically pumped magnetometers (OPMs) have made moving, wearable magnetoencephalography (MEG) possible. The OPMs typically used for MEG require a low background magnetic field to operate, which is achieved using both passive and active magnetic shielding. However, the background magnetic field is never truly zero Tesla, and so the field at each of the OPMs changes as the participant moves. This leads to position and orientation dependent changes in the measurements, which manifest as low frequency artefacts in MEG data.

**Objective:** We modelled the spatial variation in the magnetic field and used the model to predict the movement artefact found in a dataset.

**Methods:** We demonstrate a method for modelling this field with a triaxial magnetometer, then showed that we can use the same technique to predict the movement artefact in a real OPM-based MEG (OP-MEG) dataset.

**Results:** Using an 86-channel OP-MEG system, we found that this modelling method maximally reduced the power spectral density of the data by 26.2 ± 0.6 dB at 0 Hz, when applied over 5 s non-overlapping windows.

**Conclusion:** The magnetic field inside our state-of-the art magnetically shielded room can be well described by low-order spherical harmonic functions. We achieved a large reduction in movement noise when we applied this model to OP-MEG data.

**Significance:** Real-time implementation of this method could reduce passive shielding requirements for OP-MEG recording and allow the measurement of low-frequency brain activity during natural participant movement.

## I. Introduction

Magnetoencephalography (MEG) is a non-invasive functional neuroimaging technique, which can be used to localize neuronal current flow with high spatial and temporal resolution. In MEG, the magnetic field due to current flow across active neuronal populations is recorded outside of the head. At the scalp, this magnetic field is in the range of femto- to pico-Tesla [1]. These fields have typically been measured using superconducting quantum interference devices (SQUIDs). SQUID-based MEG systems consist of a large vacuum flask with a helmet shaped recess for the head that is surrounded by superconducting coils. These systems are very sensitive and have excellent dynamic range, but are stationary, expensive and require participants to remain still during the recording. Recently, compact optically pumped magnetometers (OPMs) have been developed [2]–[8]. These devices can be worn directly on the scalp and so enable participant movement during scanning [9]. Neuroscientific paradigms for MEG typically avoid any subject movement, while OPM-based MEG (OP-MEG) makes it possible to perform more naturalistic tasks [10]. Similarly, participants who struggle to remain still, such as children or people with movement disorders [11], can be more easily studied with OP-MEG.

Many varieties of OPM sensor now exist. In this work, we focus on OPMs that operate in the Spin Exchange Relaxation Free (SERF) regime, but the methods outlined below would be common to many magnetometers. One practical problem that impacts OPMs is how to maintain a fixed operating point as the participant moves. The field gradient within the OPM-dedicated Magnetically Shielded Room (MSR) at UCL is around 1000 pT/m [12]; we wish to measure fields in the femto-Tesla range (typically 0.01-1 pT). This means that any small movements of any magnetic field sensor present a considerable source of interference: 1 mm of head movement could produce a field change equivalent to a large (1 pT) brain signal. Rotations within the field cause additional artefacts.

This leads to direct and indirect artefacts from movement in OP-MEG: a direct increase or decrease in the recorded value - as described above - and a consequential change in the gain of the OPMs, which is dependent on the ambient field of the sensor [13]. The relationship between the ambient field and OPM gain means that these sensors operate optimally at magnetic fields close to zero (~± 1 nT) [14]. Movement is one common reason why the field at an OPM would step outside of this range during an OP-MEG recording. These effects usually occur at low frequency (below 4 Hz), as the movements themselves are typically low frequency (see Supplementary Fig. 1). It is partially for this reason that alpha (8–15 Hz), beta (15–30 Hz) and gamma (>30 Hz) activity has successfully been recorded with OP-MEG during movement [9], [10], [15], theta (4–8 Hz) has been recorded while the participant was unconstrained [16], but delta and infra-slow waves (<4 Hz) remain a topic of future research.

A number of methods have been suggested for minimizing changes in the background magnetic field during an OP-MEG experiment, with the most successful involving the placement of electromagnetic coils around the participant [13], [17]–[19]. The currents in these coils can be adjusted to minimize the magnetic field within the volume around the head, meaning that when the participant moves, the change in field is minimal [9], [17]. The currents in the coils can be continually updated to keep the background field close to zero, minimizing temporal changes in the background field which are introduced by external sources of interference [13], [18]. The electromagnetic coils presented in [18] have been shown to be capable of keeping the magnetic field to below 2 nT within a 40 cm × 40 cm × 40 cm box around the participant’s head, allowing the participant to move within this region. This has opened up a number of new areas of research within MEG [10], [15].

However, even when these field nulling coils are used, the magnetic field around the head is not zero. This makes rotations during OP-MEG challenging [13]. Additionally, the volume that is nulled is limited to a 40 cm box at the center of the coil system. If it were possible to record OP-MEG outside of this region, it would allow experiments that could not previously be considered; for example, recording as people walk about an MSR. This could be achieved by locally nulling the magnetic field at each sensor (using the internal coils that each sensor incorporates), as is done in some novel OPM designs that include closed-loop operating modes [20]–[24]. Here we work towards selectively controlling for movement-related field changes by creating a model of the background field. Previous simulation studies have shown how a generative model, comprised of current dipole sources located on a shell around the room, could be used to null this interference [25]. Additionally, it has been shown that modelling the background field as a spatially homogeneous mean field can offer significant improvement [26]. We explore an alternative model in which we express the low-frequency background field in the room as the sum of a real-valued spherical harmonic series [27]. Due to the wearability of OP-MEG, here we build a model of the spatial distribution of the noise from the participant’s movements. The advantage of the spherical harmonic approach is that we expect the field models to be computationally simpler to estimate and update in real-time. In this work, we establish proof-of-principle and use this information to minimize the movement-related changes in our OP-MEG recordings post-hoc.

The paper proceeds as follows. We start by mapping the magnetic field in our MSR using a triaxial magnetometer. We show how it is possible to model these field variations using relatively low-order spherical harmonic models. We then consider a typical OP-MEG experiment using the participant’s natural movements. We examine the dependence of these methods on the time-window used to construct the model. We finish by discussing the suitability of these models for correcting for movement noise in real-time.

## II. Methods

### A. Theory

A point on notation; vectors and matrices have been emboldened. Vectors are additionally italicized to differentiate the two.

For a singular OPM on the scalp, at position 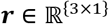 and time *t*, the recording (*B*_{*OPM*}_(***r**, t*)) is the sum of the magnetic field 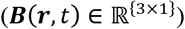 along the recording axis of the sensor 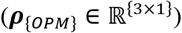 multiplied by the sensor gain (*G*), plus any sensor error terms (*e*_{*OPM*}_):

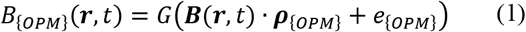

The dot indicates the dot product between the magnetic field at the sensor’s location and the orientation of its sensitive axis. ***B***(***r**, t*) has contributions from both the environment (background noise) and the brain (the signal of interest). Here we seek to model the background noise component of ***B***(***r**, t*).

We make the assumption that *G* = 1. We also assume that the sensitive axis of the sensor remains aligned with its exterior shell and that the error term, *e*_{*OPM*}_, consists of a random, Gaussian error and a static offset term. There are multiple causes of this offset, the largest being an intentionally applied field to null the initial magnetic field at the start of any experiment, described in section II.B. Additional sources of this offset include slight magnetization of the internal OPM components, effective DC fields from the cell heater, internal magnetic field gradients, and light shift (a fictitious magnetic field created from the interaction of Rubidium atoms and the laser) [28].

Equation (1) simplifies to

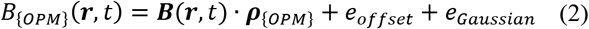

We also assume that the MSR can be approximated as a static, source-free space. The magnetic field can then be described by

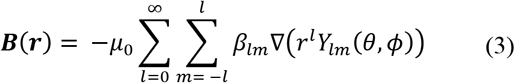

as shown by [29]–[31]. (*r,θ, ϕ*) are spherical coordinates, such that *x* = *r* sin *θ* cos *ϕ, y* = *r* sin *θ* sin *ϕ, z* = *r* cos *θ*, where *x, y* and *z* are the Cartesian coordinates. *β_lm_* are coefficients to be modelled. *Y_lm_*(*θ,ϕ*) are the spherical harmonic functions, as defined in equation (4). As the magnetic field being modelled is real, we used the real-valued – also known as solid – spherical harmonics (*S_lm_*), defined in [32] and given here in equation (5), in place of *Y_lm_* to ensure this condition. The first 6 orders are listed in Supplementary Table 1.

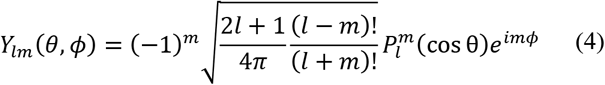

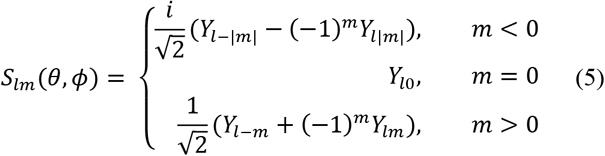

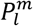 represents the associated Legendre polynomials. In equation (5), the function dependencies on *θ* and *ϕ* have been removed to keep the equation concise. Due to the nature of the associated Legendre polynomials, equation (5) can be equivalently expressed as:

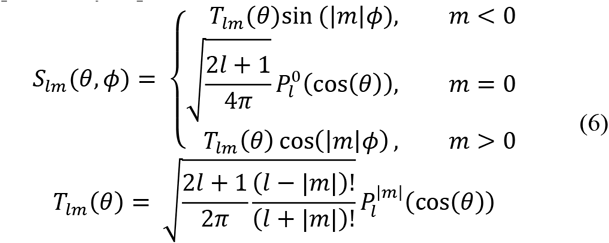

Whenever we refer to model order in this paper, we are referring to the maximum value of *l* used (*l_max_*).

We used linear regression to create the model from all recorded channels and timepoints.

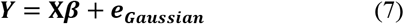

Where ***Y*** is the measured field data and **X**, the design matrix, contains the spherical harmonic model of magnetic field change over space. As such,

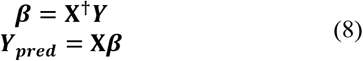

**X^†^** is the pseudoinverse of X.

The first *M* (where *M* is the number of channels) columns of the design matrix are used to account for constant (channel specific) offsets. The remaining columns describe the change in the magnetic field over space, such that the expression agrees with equations (2) and (3). The number of columns of **X** is therefore determined by the number of channels and the model complexity and is equal to (*M* + (*l_max_* + 1)^2^). The rows of X correspond to the timepoint and sensor. We chose to list all timepoints of sensor 1, then all times of sensor 2 etc. Consequently, the data for the regression (***Y***) is given as follows:

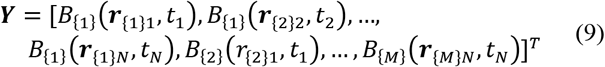

And has length (*NM*), where *N* is the number of datapoints modelled over. *B*_{*m*}_ (***r***_{*m*}*n*_, *t_n_*) refers to the recording of OPM *m* at timepoint *n* and position (***r***_{*m*}_, (*t_n_*).

As an example, in the simplest case considered here, *l_max_* = 1, and so the background magnetic field is modelled by 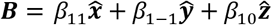, where 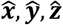 are the standard unit vectors in the direction of the *x, y* and *z* axes. In this scenario, the design matrix is as follows,

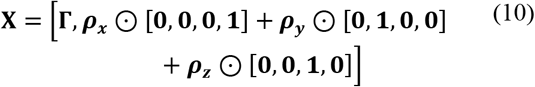

where 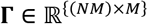 represents the matrix for the channel offsets. Each column of **Γ** corresponds to a sensor channel and is zero, unless, for column *m*, row *k*, the data in row *k* of ***Y*** was recorded by OPM channel number *m*. Apart from **Γ**, all terms in equation (10) are column vectors with values corresponding to the position and orientation of the channel in the corresponding row of ***Y***, e.g. the first element of ***ρ_x_*** is the component of the orientation of OPM channel 1 in the *x* direction at timepoint 0, the second is its orientation in *x* at timepoint 1 and so on, until the last element is the component of the orientation of OPM channel *M* in the *x* direction at timepoint *N*. **1** is a column of ones and, similarly, **0** is a column of zeros. ⊙ indicates elementwise multiplication. In this case where *l_max_* = 1, the estimated parameters would be

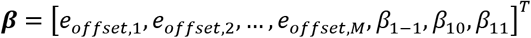

For simplicity, we have only written this out for the simplest model. To expand this to include higher order models, the real spherical harmonics, as listed in Supplementary Table 1, need only be concatenated to the inner arrays. For example, considering only the magnetic field in the *x* direction, in going from a first order model to second,

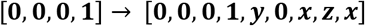

### B. Recording setup

Two experiments for two different sensor configurations – triaxial and whole-head, as described in II.C – were undertaken. Fig. 1 shows the setup of the triaxial field mapping experiment. In both cases, we used QuSpin QZFM 2^nd^ generation OPMs (https://quspin.com/products-qzfm/). The OPMs were moved manually, pseudo-randomly, either on the end of stick or on the participant’s head, around the central 1-2 cubic meters of a 4-layer MSR (Magnetic Shields, Ltd.; internal dimensions 3 m × 4 m × 2.2 m) for 5 minutes.

**Fig. 1.**
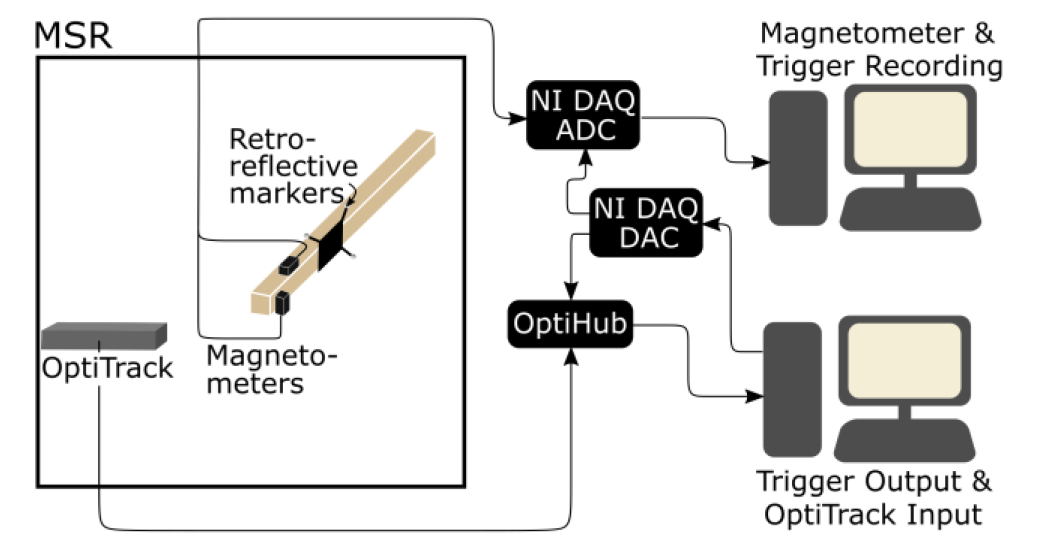
Field Mapping system set up. In the triaxial experiment, the position and orientation of two magnetometers were tracked optically, while the field along two of their axes were recorded. These two data-streams (magnetic field and position/orientation) were synchronously recorded.

Before the start of the experiments, the inner layer of mumetal lining the room was degaussed by passing a low-frequency decaying sinusoidal current through cables within the walls.

The position of a rigid array of 4 retroreflective markers, which was fixed relative to the OPMs, was recorded using an OptiTrack v120:duo motion tracking camera in the triaxial experiment or, in the case of the OP-MEG experiment, 6 OptiTrack Flex 13 cameras spaced around the room.

The magnetometer outputs were recorded at 6000 Hz using LabView with a National Instruments (NI) DAQ (NI-9205, 16-bit, ± 10 V input range), using QuSpin’s adapter (https://quspin.com/products-qzfm/ni-9205-data-acquisition-unit/). The position information was recorded on a separate computer using OptiTrack’s Motive software at 120 Hz. A 5 V voltage pulse was sent to both systems for synchronization.

Occasional occlusion of one or multiple markers led to gaps in the position data. In the triaxial recording, these gaps were filled in using cubic spline interpolation in Motive. In the OP-MEG experiment, an initial “pattern-based” interpolation was performed prior to spline fitting. In this case, when only one marker was missing (i.e. the position of the other three markers was known), the trajectory data from the other markers was used to determine the position of the occluded marker by the constraints of the rigid body. Any remaining gaps were then filled with cubic spline interpolation.

The magnetometer outputs were downsampled to 240 Hz for convenience. Linear interpolation was used to upsample the OptiTrack data to 240 Hz to match the two recordings.

The OPMs used here operate optimally in ambient magnetic fields close to zero (~ ± 1 nT), so at the start of each experiment, electromagnetic coils on board each sensor were optimized to produce a magnetic field equal and opposite to the ambient field at the time of the calibration. This bias field (typically 0.1–2 nT for our OPM-dedicated MSR) was recorded and left on throughout each recording. The gain of the OPMs was set to allow recordings of up to ± 5.56 nT. This was necessary to ensure that all of the data was within range, although, as discussed previously, larger magnitude OPM recordings have higher gain errors, meaning that there is more uncertainty in the larger field values.

The OptiTrack coordinate system was set using an initial Ground Plane recording, where a right-angled triangle frame with retroreflective markers at the corners was placed at the origin of and aligned along the axes of the desired coordinate system. This coordinate system was chosen to be related to the geometry of the MSR. The origin was set approximately in the center of the room, with the x-axis pointing towards the door, y-axis pointing down and z-axis defined so that the coordinate system is right-handed.

To find the position and orientation of the OPMs from the position of the markers, we used the Kabsch method [33] to find the optimal transformation between the known coordinates of the markers relative to the OPMs and the OptiTrack recordings. We then applied the same transformation to the known OPM coordinates and orientations at each data point.

### C. Experiments

#### 1) Triaxial Field Mapping

We created a triaxial magnetometer from two, orthogonally oriented OPMs, shown in Fig. 2. Each OPM recorded two orthogonal directions of field change, although only three axes were selected for modelling (i.e. only one of the two parallel axes was used). We recorded for 5 minutes while moving the sensor around the room, waited 20 minutes and then repeated the recording.

**Fig. 2.**
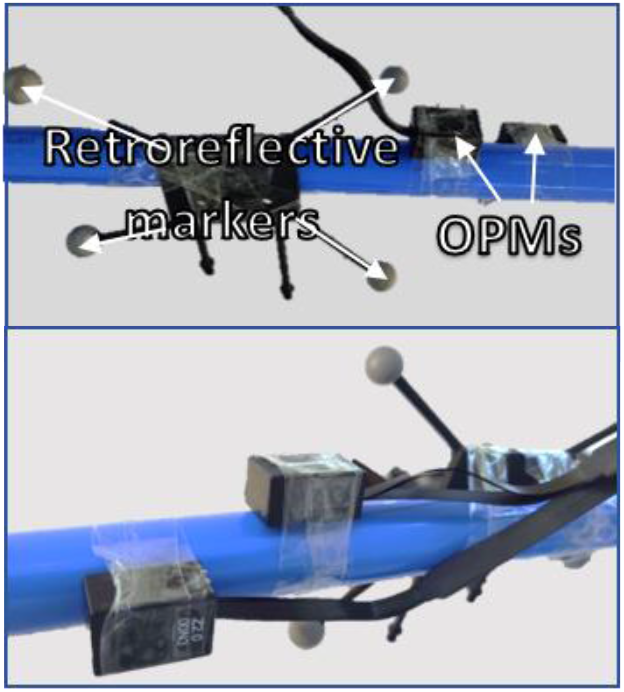
Photographs of the experimental set-up, showing the OPMs and retroreflective markers for position tracking, taken from opposite directions.

Three filters were applied to the OPM recordings: a 4^th^–order 60 Hz low-pass Butterworth filter and two 5^th^–order band-stop Butterworth filters at 50 Hz (line noise) and 120 Hz (infrared interference from the OptiTrack cameras).

To determine the number of spherical harmonic functions required to reasonably describe the magnetic field in our MSR, we tested the first six model orders. We performed a 10-fold cross-validation test to compare the different models for both recordings and evaluated their performance by the variance in the data explained by each model, quantified by the Coefficient of Determination (*R*^2^) across the full dataset.

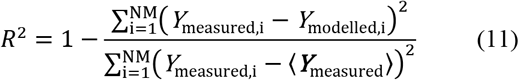

Here the brackets around ***Y***_measured_ indicate the mean over all channels and times.

Additionally, we wished to avoid overfitting and also establish whether the magnetic field changed with time. Therefore, we trained a model on the first 80% of the data and tested it on the last 20%. For this purpose, we also trained the model on the alternative run, recorded 20 minutes apart.

#### 2) OP-MEG Recording

We sought to recreate this modelling with OP-MEG data based on recordings from multiple scalp-based sensors. To create a test dataset, 43 (dual-axis) OPMs were placed evenly around a participant’s head in a 3D printed, bespoke, rigid scanner-cast. The sensor and OptiTrack marker positions relative to the scalp are shown in Fig. 3.

**Fig. 3.**
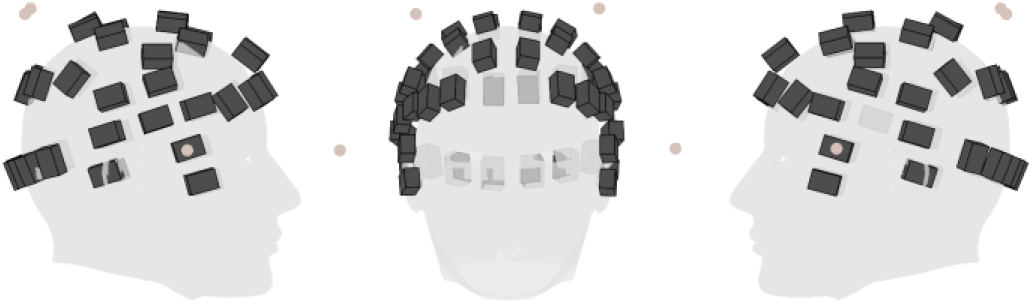
Position of each of the OPMs (black cuboids) and retroreflective markers (orange circles) in the auditory experiment. The participant’s head is represented by the grey mesh.

The participant was standing and was asked to move such that their head made large translational and rotational movements (shown in Fig. 6). The experimental protocol was approved by the UCL Research Ethics Committee and informed consent was obtained prior to participation.

Aiming to compensate for the temporal changes in the magnetic field, we performed this modelling on sliding windows of the data. Six different window lengths — 5 s, 10 s, 30 s, 60 s, 120 s and 240 s — were tested. In this work, we have consistently set the modelling step size, i.e. how often we update the model, to half of the length of the modelling window. The impact of changing this value is discussed in the supplementary material.

To evaluate the model’s performance, we looked at the shielding factor of the resulting correction, calculated with SPM (https://github.com/tierneytim/OPM).

*shielding factor (dB)*

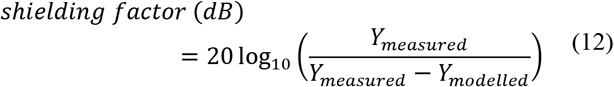

For three windows (5 s, 30 s and 120 s), we also looked at the percentage decrease in the RMS value of the OP-MEG recording and how that varied between the channels.

The position data was low-pass filtered at 2 Hz with a 6^th^– order Butterworth filter before modelling. The model predictions were low pass filtered at 2 Hz with a 5^th^–order Butterworth filter. This filter was necessary because we found that above this frequency, the noise from the motion tracking camera was larger than the noise from the movement. This is consistent with Supplementary Fig. 1, which shows that most of the movement here is described by frequency components below 2 Hz.

## III. Results

### A. Triaxial recording

The modelled magnetic field in the OPM–dedicated MSR at UCL from the whole of the first triaxial recording using a 3^rd^ order model is shown in Fig. 4. The figure shows the trajectory of the movement, which begins in the bottom left of the grid shown (as the researcher picks the sensor up from a table). The range of movement was 1.2 m, 1.4 m, and 0.8 m in *x, y*, and *z* respectively. The model is spatially smooth, as you would expect given the basis functions, with a gradient in the *x* direction (towards the door). The equivalent figures for the second triaxial run and the OP-MEG recording are shown in Supplementary Fig. 3.

**Fig. 4.**
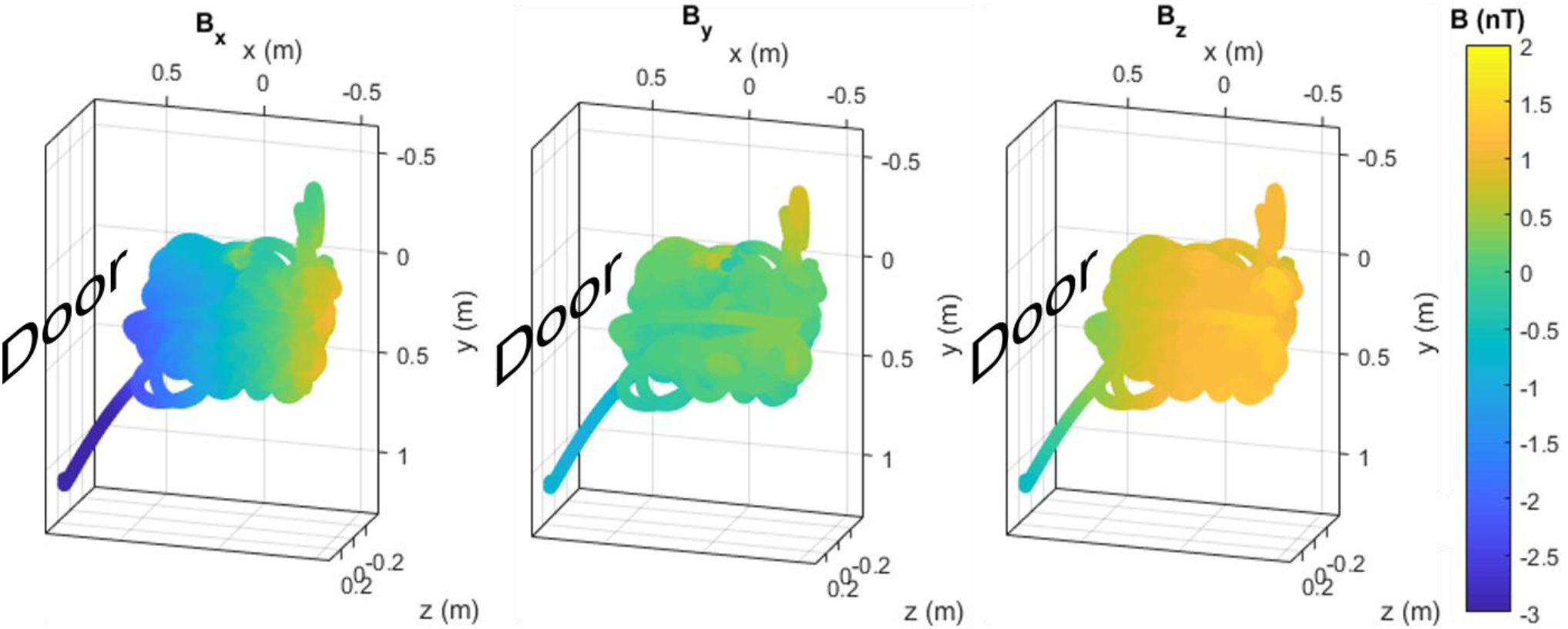
Background magnetic field in the Magnetic Shields Limited (MSL) MSR at UCL at the mean OPM position during the first run of the triaxial experiment, according to a 3rd order real spherical harmonic model. The three columns are the three magnetic field components. In each, the direction of the MSR door is indicated. The graphs are oriented to be representative of the room such that down the page is nearer to the ground in the room. The two trails coming out of the main space - bottom left and top right - are, respectively, caused by the magnetometer being picked up off the table at the start of the experiment and moving it nearer the camera (to see how this affected the field). The equivalent figure for the second run of this experiment and for the OP-MEG experiment is shown in Supplementary Fig. 4.

Fig. 5 shows the variance in both triaxial recordings for spherical harmonic models of different complexity. As one would expect, the model error decreases as the complexity of the model increases. When a 10-fold cross-validation test was performed, the difference between the within-sample variance explained and out–of–sample variance explained was within the error bars for all the models. This suggests that all the models generalize well. However, for both recordings, the same cannot be said when training on the first 80% of the data and testing on the last 20%. The model still explains over 96% of the variance in the hold out set and, indeed, the variance explained in the hold out set of the second run exceeds that in the training set but the fact that they are different implies that the magnetic field is changing in time.

**Fig. 5.**
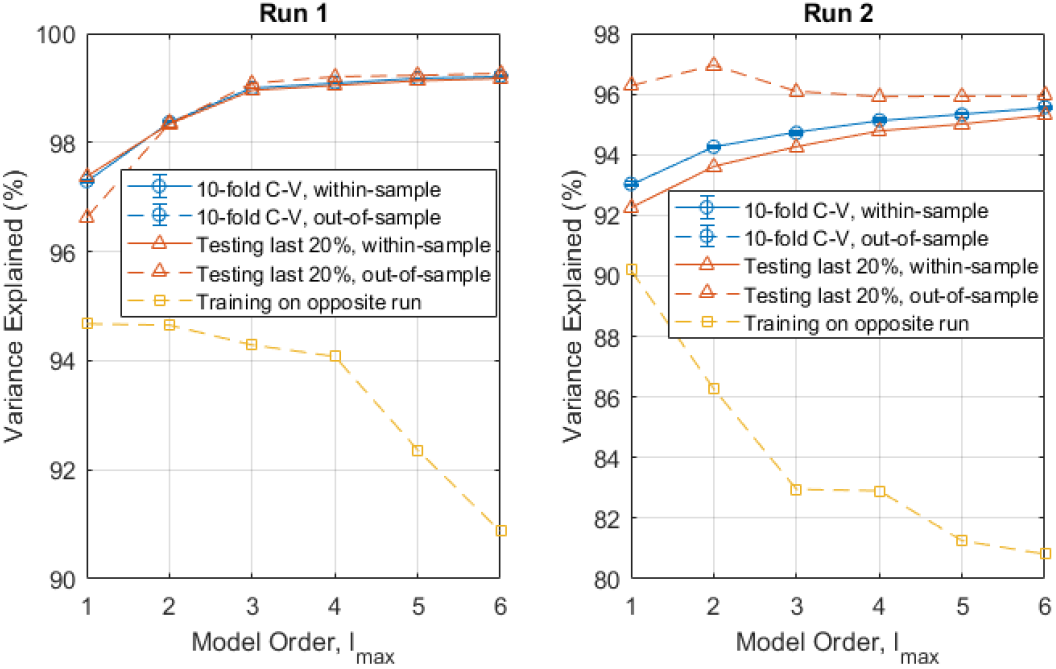
Variance explained (***R*^2^**) by different order spherical harmonic models in three different analyses: 10-fold cross-validation (blue circles), training on the first 80% of the data (orange triangles) and training on the opposite run (yellow squares). The within-sample (testing and training data are the same) variance explained is given by complete lines, the out-of-sample (testing and training data are different) variance explained is given a dashed line. The two recordings are shown on separate graphs. Run 1 (left) was recorded first, then run 2 (right) recorded 20 minutes later. Note the different scales on the two graphs.

In line with this, there is a notable drop in the variance explained when training on the alternative recording, i.e. training on data recorded in the same room, without opening or closing the door but recorded 20 minutes later or earlier. Unlike testing on the same dataset, the variance explained does not increase as the model order increases. In this case, the lower order models appear to be more stable. Additionally, the number of model parameters is (*l_max_* + 1)^2^. As the number of parameters increases, so does the size of the design matrix **X** and the time required to invert it. Therefore, for computational efficiency and robustness, a spherical harmonic model of order 2 is a pragmatic compromise for the space sampled. For the rest of the paper, if a model order is not explicitly given, a 2^nd^ order real spherical harmonic model was used.

### B. OP-MEG Recording

Having established that it is possible to describe the field in the center of the room using a low-order spherical harmonic model, we set out to examine how effective these models and estimates might be during a real participant recording. As we expect the optimal field model to change depending on the room space moved within (i.e. high orders for large spaces or spaces close to the walls), we were aiming to use the participant’s own movements to define the optimal field model, rather than reusing the models from the previous triaxial experiments.

A 100 s segment of the recordings from three randomly selected example channels, as well as the movement and rotation of the scanner-cast in the OP-MEG experiment, is shown in Fig. 6. The movements (approximately 60 cm) here are notably larger than the typical movement range for SQUID-MEG (1 cm) [34].

**Fig. 6.**
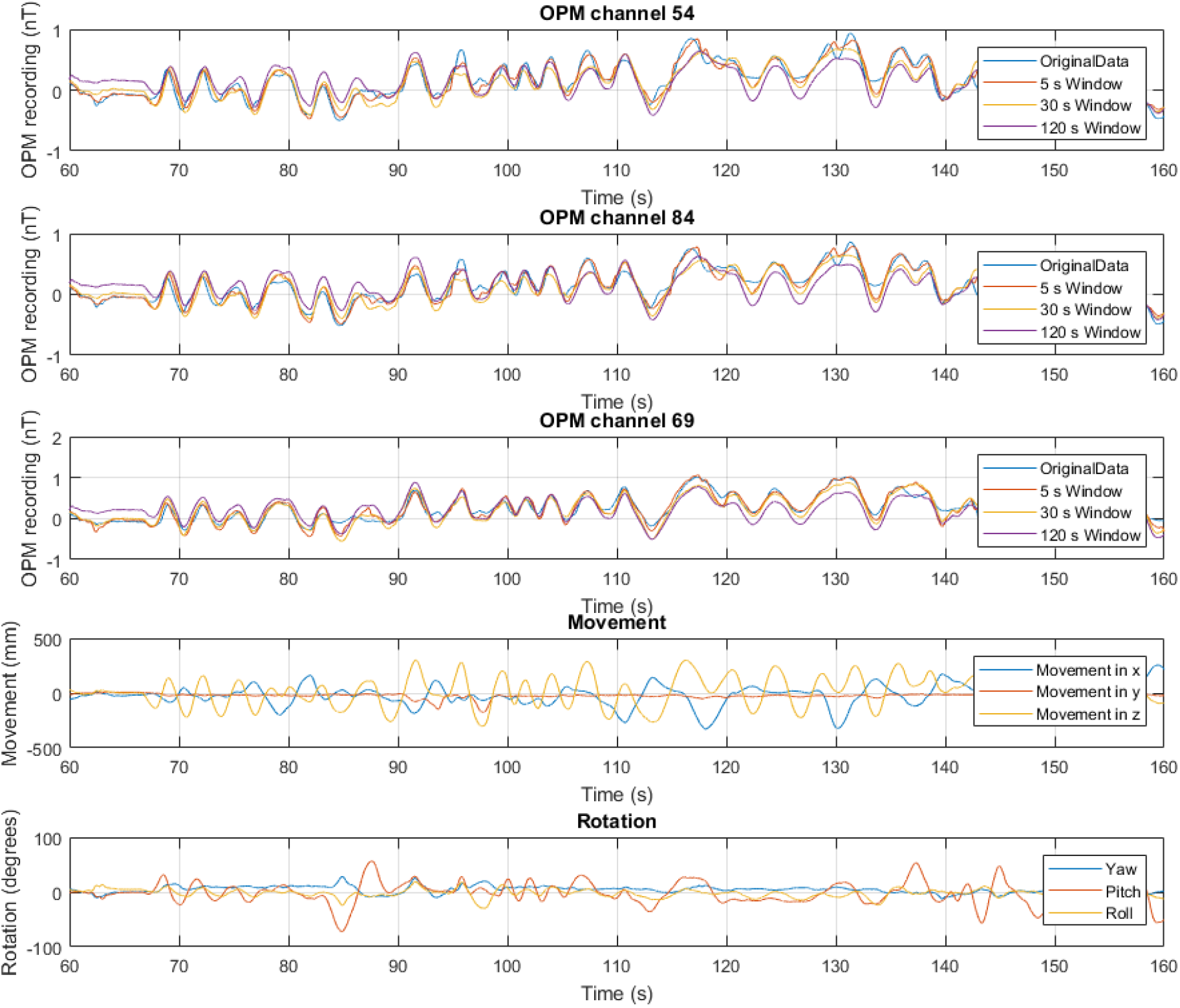
Example OPM recordings (first three rows) and corresponding movement information (last two rows) for the participant experiment. In the OPM recordings, the measured data is shown in blue. The model predictions for a first order model with three window lengths is shown: 5 s (orange), 30 s (yellow) and 120 s (purple). The position information is shown as the movement (position minus starting position, 4th row) and rotation (bottom) of the scanner-cast during the field mapping recording. In the movement panel, the x (blue), y (orange) and z (yellow) components of the position of the scanner cast in the same room-based coordinate system as the triaxial recording are shown. The bottom panel shows the pitch (blue), roll (orange) and yaw (yellow) of the scanner-cast, as recorded by the OptiTrack camera.

Fig. 6 also shows the predictions from a 2^nd^ order spherical harmonic model fit to these data. The predicted field is shown for three different sliding window lengths – 5 s, 30 s and 120 s. For each of the three channels shown, it appears from visual inspection that the 5 s window fits to the original data most closely.

The per-sample noise reduction (as defined by the percentage decrease in the root mean square of the OPM recording) for these three window lengths and all 86 channels is shown as a histogram in Fig. 7. There is variation between the 86 channels, with noise reduction value ranging from 34.9 ± 0.3 % to 66.7 ± 0.2 % for the 5 s window, with an average of 54.3 *±* 0.8 %.

**Fig. 7.**
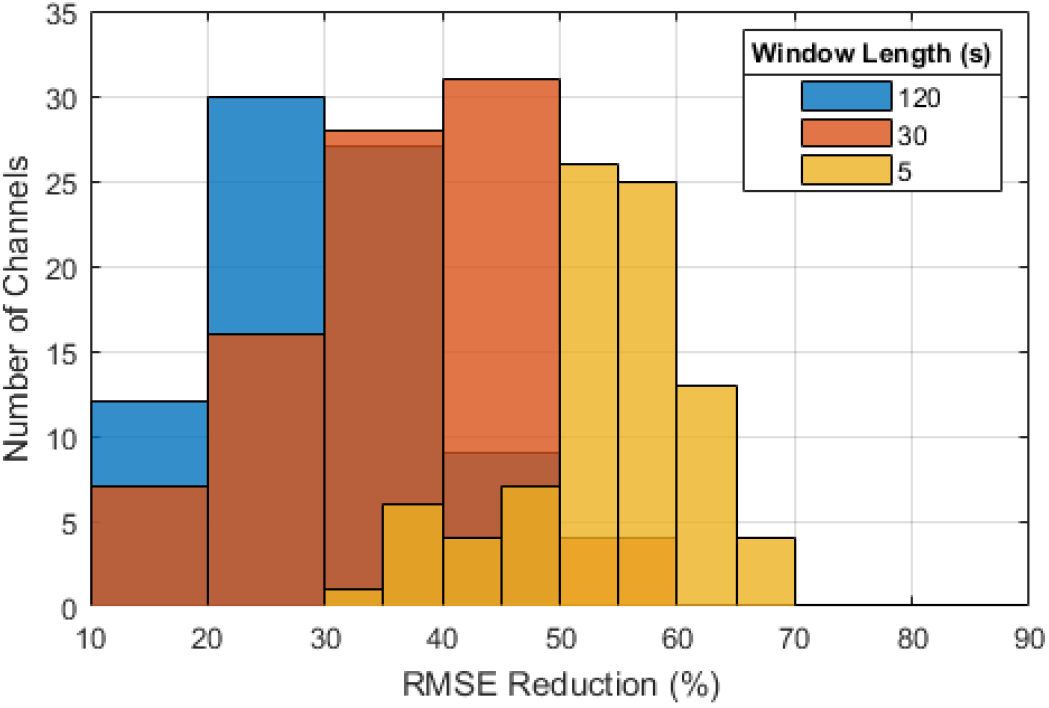
The RMS noise reduction for 120 s, 30 s and 5 s sliding modelling windows as a histogram of the values for different channels.

The level of noise reduction was found to be dependent on the length of the window used. As the window length increases, the average noise reduction decreases while the variation between channels increases. Consequently, for a 120 s window, we saw a reduction between −2.9 ± 0.5 % and 48.7 ± 0.3 % with an average of 27.1 ± 1.3 %.

To look at the dependence of performance on frequency, the shielding factor for different window lengths is examined in Fig. 8. The impact of the correction is largest at 0 Hz for all window lengths, with maxima at 7.8 ± 0.5 dB, 13.3 ± 0.4 dB and 26.2 ± 0.6 dB for 120 s, 30 s and 5 s windows respectively. However, above 1 Hz, particularly for the 5 s and 10 s windows, the algorithm can have a detrimental effect, with shielding factors below 0 dB suggesting that the additional noise from the OptiTrack introduced by applying the correction is higher than the original movement noise in this region.

**Fig. 8.**
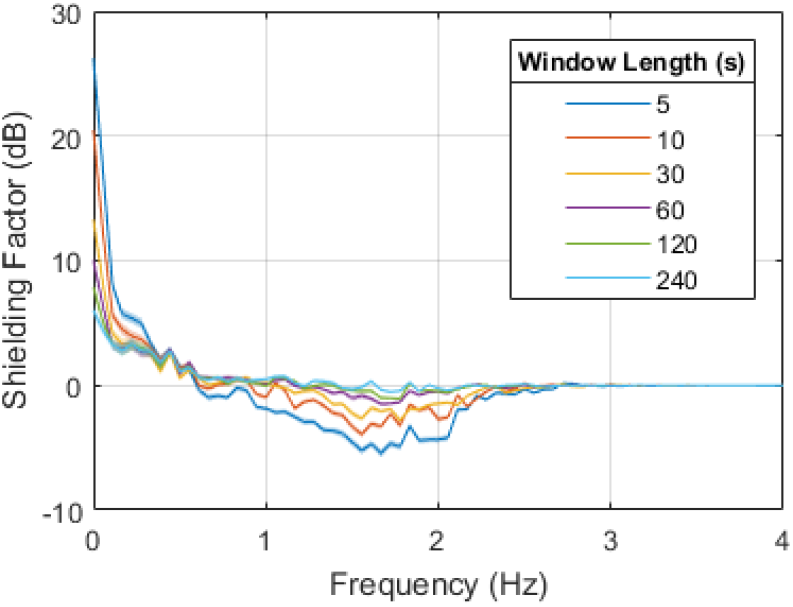
Shielding factor for a 2^nd^ order spherical harmonic model on the OP-MEG recording for different window lengths. The values shown are the mean over all channels, with the width of the line given by the standard error of the mean.

When longer (60 s, 120 s and 240 s) windows are used, the algorithm is less detrimental; the noise in the position recordings has less of an impact by simply having more datapoints from which to create the model. This is also the case for the out-of-sample shielding factor, shown in Supplementary Fig. 5. However, in this scenario, the field modelling also has limited benefit, with a maximum shielding factor of 8.9 *±* 0.7 dB for the 120 s window. Along with model complexity, modelling window length will be an important factor to be considered when using this method to reduce movement noise in OP-MEG.

## IV. Discussion

We tested a method to compensate for sensor movement within the central portion of a magnetically shielded room using a spherical harmonic field model. We created models from recordings made while moving a triaxial sensor and a whole-head sensor array. We showed that low-order spherical harmonics could explain (and predict) over 80 % of the variance in the data.

We used the same spherical harmonic models with an on-scalp array but we note the performance gains were much less striking. Although large noise reduction was achieved for short time windows (54.3 ± 0.8 % at 5 s) with the on-scalp array, the performance for longer windows was relatively modest. This is likely due to a number of factors. First, the magnetic field in the room was changing temporally as well as spatially, due, for example, to passing traffic. Second, there was additional noise due to movement of the sensor cabling. These cables pull on and consequently move the magnetically sensitive cell within the OPM housing, creating field changes due to internal device movement. The cables also interact as they move across one another, creating movement-related but unpredictable artefacts. These issues are currently being resolved with improved cable fastening and layout. These factors may help to explain the poorer performance at long window lengths and why, when we repeated the 10-fold cross-validation used in the triaxial experiment on the OP-MEG data, the variance explained was notably lower (see Supplementary Fig. 3). One future improvement to the method could be to add regularization, in particular on the OPM offsets which should change far more slowly than the background magnetic field. A Bayesian update method for example would allow some parameters to be updated more slowly than others.

This method draws inspiration from the signal space separation method (SSS) [30], [35], [36] and mean field modelling of the background magnetic field in an MSR [26]. SSS makes use of the spherical harmonic description of the magnetic field to separate fields arising from within and outside of the head. Here we make use of the fact that the head is moving and assume that brain activity is negligible compared to the movement induced artefacts. The assumption in mean field modelling is that the background magnetic field across the head is spatially constant and can therefore be removed. This takes place time-point by time-point without any knowledge of head-position. It is therefore well-suited to temporally non-stationary interference. In contrast, here we assume that the background magnetic field varies spatially and only changes slowly in time. This makes the model slower to compute but, critically, means that the magnetic field at a new position, orientation, and time can be predicted. This may have advantages in situations in which movements are fast or where field gradients are high, and field changes would have otherwise moved the OPMs outside of their optimal operating range (see next section). There is clearly scope for further work in which all three approaches are combined.

Here we discussed the change in the OPM recordings from movement while the OPMs are operating within their dynamic range of ±5.56 nT [19]. We are fortunate that the central part of our room meets these specifications after degaussing the inner mu-metal layer, but for other rooms or different ranges of movement, we envisage that real-time field correction may be required to keep the OPMs within their operating range. If the sensors can continuously be kept close to their optimal (0 T) operating point this also mitigates the gain errors (~1 % per nT [9], [13], [14], [37]) which are incurred as a result of operating at an offset field during movement. Practical constraints to be considered in the future would be the cross-talk from these compensating fields between the coils on the different OPMs [38]. The space the participant moves through will also likely be important in the choice of model parameters, in particular the window length and model order. Here we looked at continuous, large movements; arguably the worst-case-scenario for OP-MEG recordings. However, typical neuroimaging experiments are likely to contain less frequent and smaller movements. A longer modelling window and a lower order model may be preferable in these situations.

The timing of the applied field will also likely be important. In the way we have used field modelling in this paper, time is not a significant limitation, since the modelling is done offline after an experiment. However, in this real-time scenario, computation time will be critical and should not be more than the time between recalculating the model. This recalculation time will depend to some extent on the stationarity of the environment. It is encouraging, therefore, that in Fig. 5, over 80% of the field variation in the room can be predicted from measurements taken 20 minutes apart. The computation time is dependent on the size of the design matrix, itself determined by the number of OPM channels, number of datapoints in a modelling window, and model complexity. The time between recalculating the model, equivalent to the step size for the sliding window, should be chosen to be small enough to account for the changes in magnetic field with time, but larger than the computation time. In this paper, we have consistently used a step size of half the sliding window length. The relationship between this step size and the model accuracy, computation time, and noise reduction is discussed in the supplementary material. One of the advantages of a spherical harmonic model as we have used here, is the relative simplicity by comparison with a more complicated source model [25]. Further work will be needed to increase the speed of this modelling, in order to selectively correct for movement-related field changes in real-time.

We have focused on mapping signal modulations due to movements through static fields; we are currently exploring how this method could be applied to predict spatial variations in other interfering fields. For example, the mains (50 Hz) noise within our shielded room has a clear spatial structure. Other fields, such as those due to the vibration of the room walls may also have a deterministic structure. Typically, one might use an adaptive filter and reference noise measurements to minimize such signals; however, updating the correlation between the reference sensor and signal recordings over time as the participant moves introduces high pass filtering effects. Using a method based purely on measurement of spatial position would incur no sacrifice in recording bandwidth as the optimal weighting of the reference signal(s) would be given by the location and orientation of each sensor.

## V. Conclusion

In summary, we have explored a method to model the spatial variation of background magnetic fields within a magnetically shielded room and the interference they cause during an OP-MEG recording. We used a spherical harmonic field model and found that for the central portion of our shielded room, effective field cancellation could be achieved for low model orders. This is consistent with prior work and encouraging for future use of external field nulling coils typically used in OP-MEG, which are generally capable of producing 1^st^ order magnetic field gradients [13], [17], [18].

This demonstrates the potential for real-time correction based on these models in the future. These preliminary steps hold promise for OP-MEG systems with greater movement tolerance requiring less passive shielding.

## Acknowledgements

This work was supported by a Wellcome collaborative award to GRB, MJB and RB (203257/Z/16/Z, 203257/B/16/Z), the EPSRC-funded UCL Centre for Doctoral Training in Medical Imaging (EP/L016478/1), the Department of Health’s NIHR-funded Biomedical Research Centre at University College London Hospitals and EPSRC (EP/T001046/1) funding from the Quantun Technology hub in sensing and timing (sub-award QTPRF02). NA, RAS and EAM are supported by a Wellcome Principal Research Fellowship to EAM (210567/Z/18/Z). The Wellcome Centre for Human Neuroimaging is supported by core funding from the Wellcome [203147/Z/16/Z]. The authors would also like to thank Vishal Shah and QuSpin, and Mark Lim and Chalk Studios for their continued support.

