## Supplementary Material for "Magnetic Field Mapping and Correction for Moving OP-MEG"

### I. SUPPLEMENTARY FIGURES

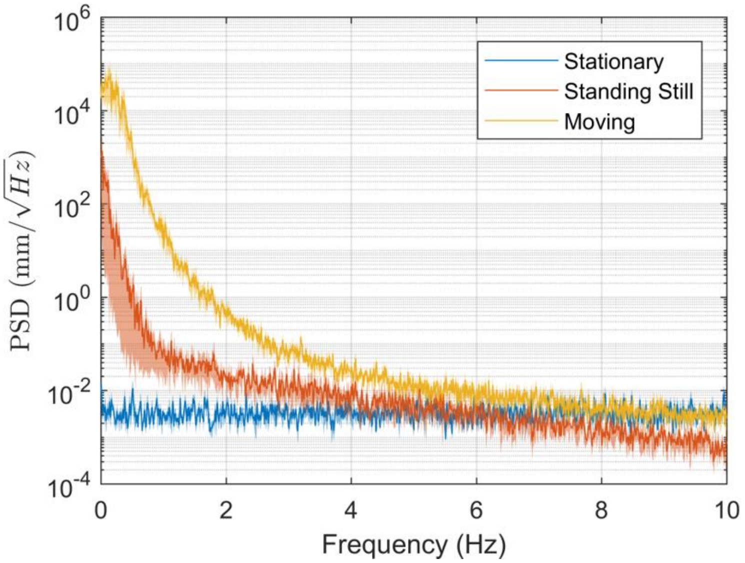


Supplementary Fig. 1. Power spectral density (PSD) of the participant movement, as recorded with OptiTrack Flex 13 cameras, in the OP-MEG experiment while asked to move (yellow), in a separate recording where they stood as still as possible (orange) and in a recording with the scanner-cast sitting stationary on the table (blue). Each recording was 5 minutes long. The position data was demeaned before the PSD was estimated using Welch's method, with 100s segments. The figure shows the average over the three dimensions. The width of the line is given by the standard error of the mean.

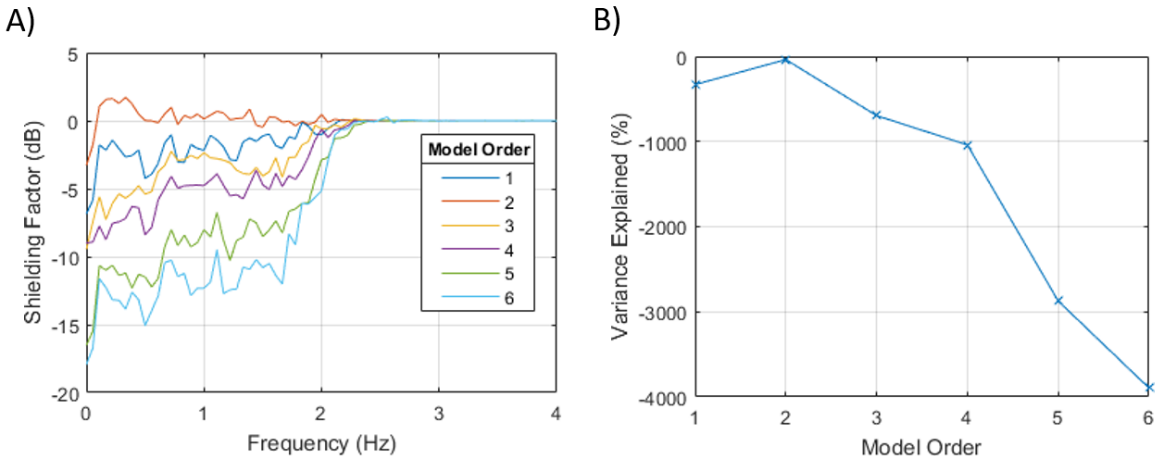


Supplementary Fig. 2. Noise correction when the model was trained on triaxial data run 1 and then used to predict the data from the on-scalp array . Evaluated by (A) the shielding factor and (B) the variance explained. There are two likely explanations for the poor performance: 1) the coordinate system of the two recordings is not perfectly aligned or 2) the magnetic field in the room changes with time, particularly as items are added and removed from the room and the door is opened and closed, magnetizing the mu-metal of the room.

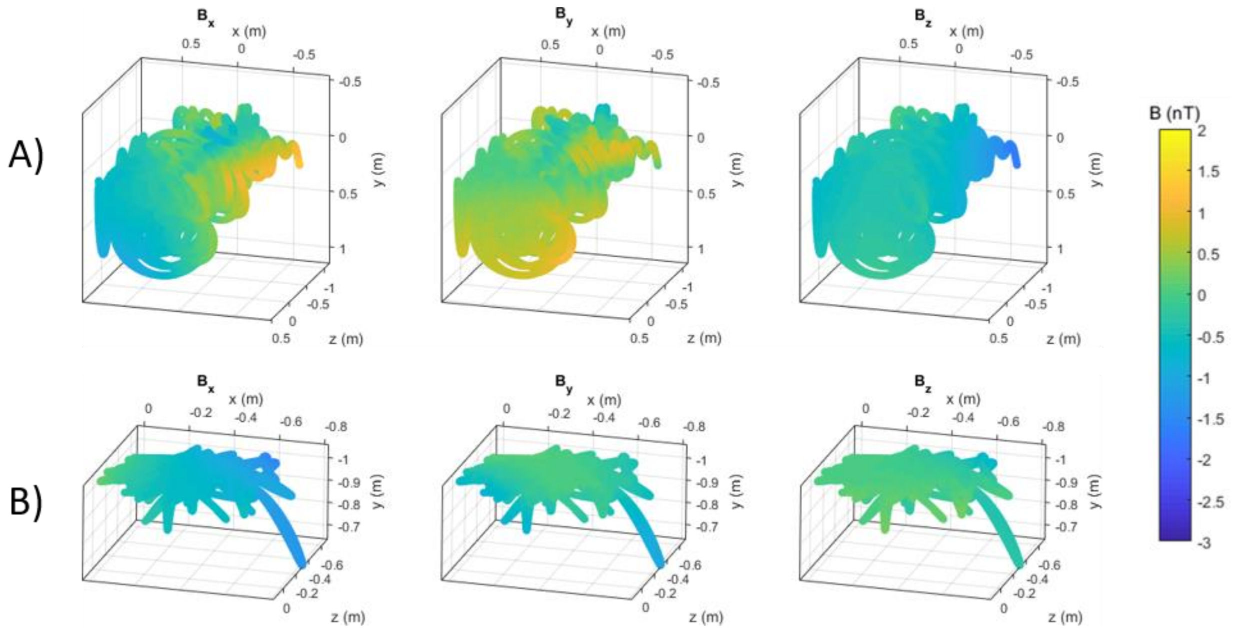


Supplementary Fig. 4. Modelled (using a 3rd order real spherical harmonic model) magnetic field at the mean OPM position during (A) run 2 of the triaxial experiment, (B) the participant on-scalp experiment. The three columns are the three magnetic field components.

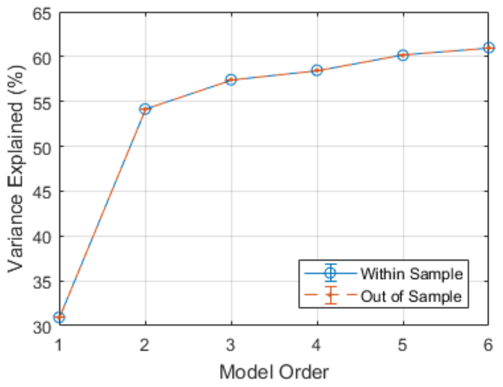


Supplementary Fig. 3. Variance explained ($\boldsymbol{R^2}$) by different order spherical harmonic models in the participant data, from a 10-fold cross-validation analysis. The blue circles are the within-sample variance explained, the orange dots are the out-of-sample variance explained.

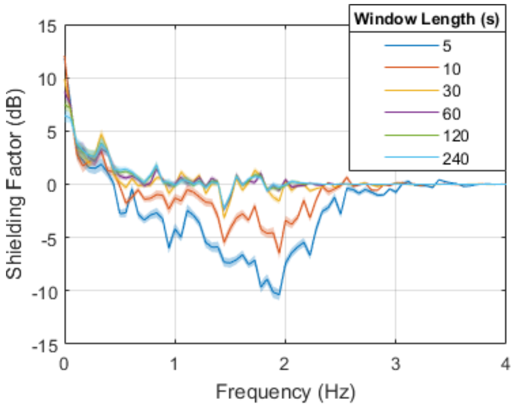


Supplementary Fig. 5. Out-of-sample Shielding Factor for different window lengths in the OP-MEG experiment. For the last 60 s of the recording, the background magnetic field was predicted one second after a window. Only the last 60 s of the recording was used to allow fair comparison between the different windows, since it was a 300 s recording. The line shown is the mean across channels; the width of the line is the standard error of the mean.

#### II. Supplementary Table

| $l$ | $m$ | $r^l S_{lm}$ | $\frac{\partial(r^l S_{lm})}{\partial x}$ | $\frac{\partial(r^l S_{lm})}{\partial y}$ | $\frac{\partial(r^l S_{lm})}{\partial z}$ |
|---|---|---|---|---|---|
| 0 | 0 | 1 | 0 | 0 | 0 |
| | -1 | $y$ | 0 | 1 | 0 |
| 1 | 0 | $z$ | 0 | 0 | 1 |
| | 1 | $x$ | 1 | 0 | 0 |
| | -2 | $xy$ | $y$ | $x$ | 0 |
| | -1 | $yz$ | 0 | $z$ | $y$ |
| 2 | 0 | $2z^2 - x^2 - y^2$ | $x$ | $y$ | $z$ |
| | 1 | $zx$ | $z$ | 0 | $x$ |
| | 2 | $x^2 - y^2$ | $x$ | $y$ | 0 |
| | -3 | $(3x^2 - y^2)y$ | $xy$ | $x^2 - y^2$ | 0 |
| | -2 | $xyz$ | $yz$ | $xz$ | $xy$ |
| | -1 | $y(x^2 + y^2 - 4z^2)$ | $xy$ | $x^2 + 3y^2 - 4z^2$ | $zy$ |
| 3 | 0 | $z(2z^2 - 3x^2 - 3y^2)$ | $xz$ | $yz$ | $6z^2 - 3x^2 - 2y^2$ |
| | 1 | $x(x^2 + y^2 - 4z^2)$ | $2x^2 + y^2 - 4z^2$ | $xy$ | $xz$ |
| | 2 | $(x^2 - y^2)z$ | $xz$ | $yz$ | $x^2 - y^2$ |
| | 3 | $(3y^2 - x^2)x$ | $x^2 - y^2$ | $xy$ | 0 |
| | -4 | $xy(x^2 - y^2)$ | $y(3x^2 - y^2)$ | $x(x^2 - 3y^2)$ | 0 |
| | -3 | $(3x^2 - y^2)yz$ | $xyz$ | $z(x^2 - y^2)$ | $y(3x^2 - y^2)$ |
| | -2 | $xy(7z^2 - r^2)$ | $y(3x^2 + y^2 - 6z^2)$ | $x(x^2 + 3y^2 - 6z^2)$ | $xyz$ |
| | -1 | $yz(yz^2 - 3r^2)$ | $xyz$ | $z(3x^2 + 9y^2 - 4z^2)$ | $y(x^2 + y^2 - 4z^2)$ |
| 4 | 0 | $35z^4 - 30z^2r^2 + 3r^4$ | $x(x^2 + y^2 - 4z^2)$ | $y(x^2 + y^2 - 4z^2)$ | $z(3x^2 + 3y^2 - 2z^2)$ |
| | 1 | $xz(7z^2 - 3r^2)$ | $z(9x^2 + 3y^2 - 4z^2)$ | $xyz$ | $x(x^2 + y^2 - 4z^2)$ |
| | 2 | $(x^2 - y^2)(7z^2 - r^2)$ | $x(x^2 - 3z^2)$ | $y(y^2 - 3z^2)$ | $z(x^2 - y^2)$ |
| | 3 | $(x^2 - 3y^2)xz$ | $z(x^2 - y^2)$ | $xyz$ | $3xy^2 - x^3$ |
| | 4 | $x^4 - 6x^2y^2 + y^4$ | $x(x^2 - 3y^2)$ | $y(y^3 - 3x^2)$ | 0 |
| | -5 | $y(5x^4 - 10x^2y^2 + y^4)$ | $xy(x^2 - y^2)$ | $x^4 - 6x^2y^2 + y^4$ | 0 |
| | -4 | $xyz(x^2 - y^2)$ | $yz(3x^2 - y^2)$ | $xz(x^2 - 3y^2)$ | $xy(x^2 - y^2)$ |
| | -3 | $y(3x^2 - y^2)(r^2 - 9z^2)$ | $xy(3x^2 + y^2 - 12z^2)$ | $3x^4 + 6x^2y^2 - 24x^2z^2 - 5y^4 + 24y^2z^2$ | $yz(3x^2 - y^2)$ |
| | -2 | $xyz(x^2 + y^2 - 2z^2)$ | $yz(3x^2 + y^2 - 2z^2)$ | $xz(x^2 + 3y^2 - 2z^2)$ | $xy(x^2 + y^2 - 6z^2)$ |
| | -1 | $y(x^4 + 2x^2y^2 - 12x^2z^2 + y^4 - 12y^2z^2 + 8z^4)$ | $xy(x^2 + y^2 - 6z^2)$ | $x^4 + 6x^2y^2 - 12x^2z^2 + 5y^4 - 36y^2z^2 + 8z^4$ | $yz(3x^2 + 3y^2 - 4z^2)$ |
| 5 | 0 | $63z^5 + 15zr^4 - 70z^3r^2$ | $xz(3x^2 + 3y^2 - 4z^2)$ | $yz(3x^2 + 3y^2 - 4z^2)$ | $3r^4 + 35z^4 - 30z^2r^2$ |
| | 1 | $x(x^4 + 2x^2y^2 - 12x^2z^2 + y^4 - 12y^2z^2 + 8z^4)$ | $5x^4 + 6x^2y^2 - 36x^2z^2 + y^4 - 12y^2z^2 + 8z^4$ | $xy(x^2 + y^2 - 6z^2)$ | $xz(3x^2 + 3y^2 - 4z^2)$ |
| | 2 | $z(x^2 - y^2)(x^2 + y^2 - 2z^2)$ | $xz(x^2 - z^2)$ | $yz(y^2 - z^2)$ | $(x^2 - y^2)(x^2 + y^2 - 6z^2)$ |
| | 3 | $(3xy^2 - x^3)(r^2 - 9z^2)$ | $3(x^2 - y^2)(r^2 - 9z^2) - 2x^2(3y^2 - x^2)$ | $xy(x^2 + 3y^2 - 12z^2)$ | $z(3xy^2 - x^3)$ |
| | 4 | $z(x^4 - 6x^2y^2 + y^4)$ | $xz(x^2 - 3y^2)$ | $yz(3x^2 - y^2)$ | $x^4 - 6x^2y^2 + y^4$ |
| | 5 | $x^5 - 10x^3y^2 + 5xy^4$ | $x^4 - 6x^2y^2 + y^4$ | $xy(x^2 - y^2)$ | 0 |
| | -6 | $xy(3x^4 - 10x^2y^2 + 3y^4)$ | $y(5x^4 - 10x^2y^2 + y^4)$ | $x(3x^4 - 30x^2y^2 + 15y^4)$ | 0 |
| | -5 | $yz(5x^4 - 10x^2y^2 + y^4)$ | $xyz(x^2 - y^2)$ | $z(5x^4 - 30x^2y^2 + 5y^4)$ | $y(5x^4 - 10x^2y^2 + y^4)$ |
| | -4 | $xy(x^2 - y^2)(x^2 + y^2 - 10z^2)$ | $y(-x^4 + 6x^2z^2 - 2y^2z^2)$ | $x(-x^4 + 10x^2z^2 + 5y^4 - 30y^2z^2)$ | $xyz(x^2 - y^2)$ |
| | -3 | $yz(3x^2 - y^2)(3x^2 + 3y^2 - 8z^2)$ | $xyz(3x^2 + y^2 - 4z^2)$ | $z(3x^4 + 6x^2y^2 - 8x^2z^2 - 5y^4 + 8y^2z^2)$ | $y(3x^2 - y^2)(x^2 + y^2 - 8z^2)$ |
| | -2 | $xy(x^4 + 2x^2y^2 - 16x^2z^2 + y^4 - 16y^2z^2 + 16z^4)$ | $y(5x^4 + 6x^2y^2 - 48x^2z^2 + y^4 - 16y^2z^2 + 16z^4)$ | $x(x^4 + 6x^2y^2 - 16x^2z^2 + 5y^4 - 48y^2z^2 + 16z^4)$ | $xyz(x^2 + y^2 - 2z^2)$ |
| | -1 | $yz(5x^4 + 10x^2y^2 - 20x^2z^2 + 5y^4 - 20y^2z^2 + 8z^4)$ | $xyz(x^2 + y^2 - 2z^2)$ | $z(5x^4 + 30x^2y^2 - 20x^2z^2 + 25y^4 - 60y^2z^2 + 8z^4)$ | $y(x^4 + 2x^2y^2 - 12x^2z^2 + y^4 - 12y^2z^2 + 8z^4)$ |
| 6 | 0 | $5r^6 + 15x^2y^2(x^2 + y^2) - 90z^2r^4 - 180x^2y^2z^2 + 120z^4r^2 - 51z^6$ | $x(x^4 + 2x^2y^2 - 12x^2z^2 + y^4 - 12y^2z^2 + 8z^4)$ | $y(x^4 + 2x^2y^2 - 12x^2z^2 + y^4 - 12y^2z^2 + 8z^4)$ | $z(15x^4 + 30x^2y^2 - 40x^2z^2 + 15y^4 - 40y^2z^2 + 8z^4)$ |
| | 1 | $xz(5r^4 + 10x^2y^2 - 20x^2z^2 - 20y^2z^2 + 3z^4)$ | $z(25x^4 + 30x^2y^2 - 60x^2z^2 + 5y^4 - 20y^2z^2 + 8z^4)$ | $xyz(x^2 + y^2 - 2z^2)$ | $x(x^4 + 2x^2y^2 - 12x^2z^2 + y^4 - 12y^2z^2 + 8z^4)$ |
| | 2 | $(x^2 - y^2)(r^4 + 2x^2y^2 - 16r^2z^2 + 31z^4)$ | $x(3x^4 + 2x^2y^2 - 32x^2z^2 - y^4 + 16z^4)$ | $y(-x^4 + 2x^2y^2 + 3y^4 - 32y^2z^2 + 16z^4)$ | $z(x^2 - y^2)(x^2 + y^2 - 2z^2)$ |
| | 3 | $z(3xy^2 - x^3)(3x^2 + 3y^2 - 8z^2)$ | $z(-5x^4 + 6x^2y^2 + 8x^2z^2 + 3y^4 - 8y^2z^2)$ | $xyz(x^2 + 3y^2 - 4z^2)$ | $x(x^2 - 3y^2)(r^2 - 9z^2)$ |
| | 4 | $(x^4 - 6x^2y^2 + y^4)(x^2 + y^2 - 10z^2)$ | $x(-3x^4 + 10x^2y^2 + 20x^2z^2 + 5y^4 - 60y^2z^2)$ | $y(5x^4 + 10x^2y^2 - 60x^2z^2 - 3y^4 + 20y^2z^2)$ | $z(x^4 - 6x^2y^2 + y^4)$ |
| | 5 | $z(x^5 - 10x^3y^2 + 5xy^4)$ | $z(5x^4 - 30x^2y^2 + 5y^4)$ | $xyz(x^2 - y^2)$ | $x^5 - 10x^3y^2 + 5xy^4$ |
| | 6 | $x^6 - 15x^2y^2(x^2 - y^2) - y^6$ | $x(x^4 - 10x^2y^2 + 5y^4)$ | $y(5x^4 - 10x^2y^2 + y^4)$ | 0 |

Supplementary Table 1. Real-valued spherical harmonic functions for the first 6 orders, multiplied by $\boldsymbol{r^l}$, and their partial derivatives with respect to $\boldsymbol{x}$, $\boldsymbol{y}$ and $\boldsymbol{z}$. $\boldsymbol{x}$, $\boldsymbol{y}$ and $\boldsymbol{z}$ are the three displacement ($\boldsymbol{r}$) components and $\boldsymbol{r} = |\boldsymbol{r}| = \sqrt{\boldsymbol{x^2 + y^2 + z^2}}$. All multiplying constants have been ignored for convenience.

#### III. Other Supplementary Material

##### 1) Modelling window step size and timing considerations

In this paper, when modelling over windows of the data (as opposed to the entirety of the dataset), we used a sliding window. We slid the window by steps of half the window size. In other words, for a 10 s window, we modelled on 10 s of data and used that model to predict the last 5 s of the original 10, then moved the window forward 5 s and modelled on the next 10 s. Here we look at the impact of changing that step size.

*Methods*

We repeated the analysis of the participant data, this time keeping the window length fixed at 10 s and changing the step size. We only used the first 20 s of data and used a 1st order model rather than 2nd to minimize the time to run the experiment. We looked at the median (across channels) shielding factor, root mean (over all data) square error or the residuals and total computation time.

*Results*

Supplementary Fig. 6 shows that the model performance was not greatly impacted by increasing or reducing the step size. If anything, increasing the step size improved the performance, with the RMSE (over all time and channels) at a minimum at half the window length, as we have used throughout the paper. The computation time is greater when a smaller step size is used however, since the model needs to be recalculated more often.

We do observe that the computation time per model calculation increases with the step size. This is understandable, since while the size of the window we are modelling on remains the same, the number of datapoints we calculate at the end is smaller when the step size is smaller.

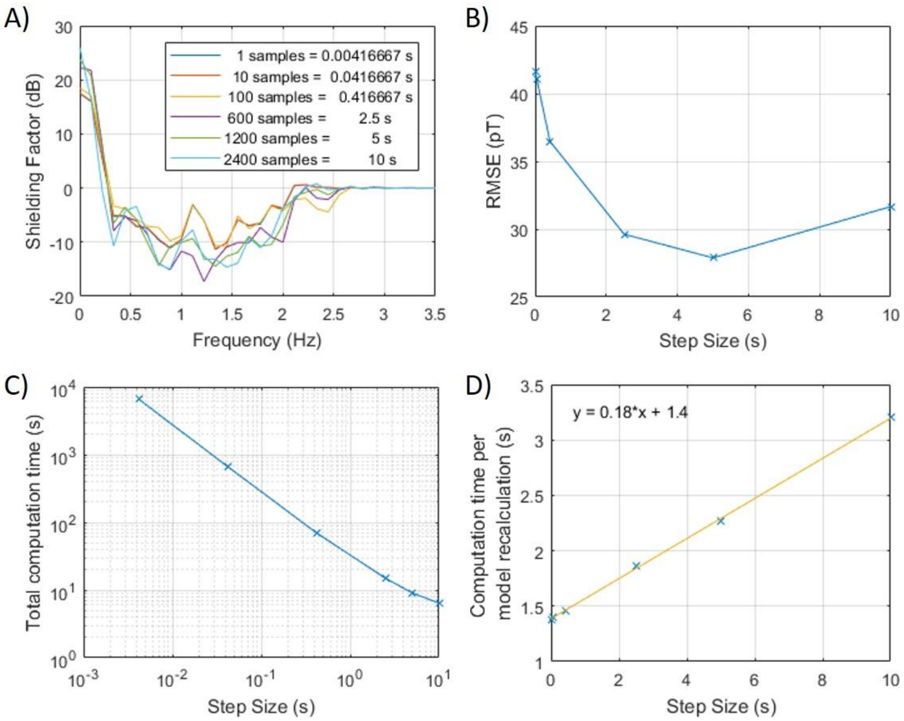


Supplementary Fig. 6 Impact of changing the modelling step size. A) shows the shielding factor for the different modelling step sizes tested. There does not appear to be a difference between the performance, apart from at near DC frequencies where higher step sizes seem to give an advantage. This is reflected in B), which shows the RMSE across the data. C) shows the total computation time while D) shows C) divided by the number of times the model is recalculated, i.e. the computation time per model calculation. D) has been linearly fitted using Matlab's fitting toolbox. The equation for the line is shown in the top left corner of the graph.

*Discussion*

Ideally, we would like to recalculate the model at every datapoint, but these results demonstrate why that is unreasonable. While there is no firm time constraint on this modelling, 6612 s for 20 s of data (1/15 of the whole dataset) is clearly untenable. It is for this reason that we have used half the window length (in this case 5 s or 1200 datapoints) in this paper.

Since we are looking to use this model to update the magnetic field at the OPMs in real time, computation time will become a significant consideration. For this to work, we require the time to compute the model to be less than the step size. For this 10 s window, these results imply that this occurs for a 1.71 s, or 410 samples (since the sample rate was 240 Hz), step size and longer.
